# High-throughput screening reveals diverse lambdoid prophage inducers

**DOI:** 10.1101/2025.05.29.656831

**Authors:** Gayatri Nair, Rabia Fatima, Alexander. P. Hynes

## Abstract

Prophages; dormant bacteriophage genomes integrated within the bacterial chromosome, play pivotal roles in shaping microbial communities when awakened. Our current understanding of prophage activation is largely shaped by a narrow set of traditional DNA-damaging inducers, such as mitomycin C and ciprofloxacin, which trigger the bacterial SOS response. This study employed high-throughput screening of 3,921 compounds to identify novel prophage inducers using model lambdoid prophage HK97. We identified multiple new inducers across diverse pharmacological classes, including dietary supplements and therapeutics. Despite the variety in compounds, all acted through SOS-dependent pathways. However, bleomycin, an antineoplastic antibiotic, demonstrated broad-spectrum and potent prophage induction exceeding standard inducers, with activity validated across multiple phages and bacterial hosts. These findings expand the repertoire of prophage inducers into commonly ingested xenobiotics, and introduce Bleomycin as a powerful, cost-effective tool for prophage research.

## Introduction

Bacteriophages (phages), the most abundant biological entities on Earth, have their genomes integrated as prophages in 75% of sequenced bacterial genomes (1, 2). Far from being inert, these prophages can profoundly alter bacterial behavior and shape microbial ecosystems when “awakened” (3-5). Despite their prevalence and impact, our understanding of prophage activation – known as induction - has been shaped by a handful of experimental tools. Most studies rely on synthetic agents like Mitomycin C, and Ciprofloxacin, as well as Ultraviolet (UV) light, to stimulate prophage induction in model systems like lambda, a temperate phage of *Escherichia coli* (6-8). Exposure to these strongly and reproducibly induces the bacterial SOS response, which in turn triggers prophage induction (8-10). While effective, this approach has become the default across studies, potentially biasing our view of phage–host interactions and limiting the discovery of alternative inducers that may be more relevant to natural environments or clinical contexts.

Compared to other environments, the human microbiome is especially enriched for temperate phages, which predominantly exist in a lysogenic state by integrating their genomes into the host chromosome and persisting as prophages (11-13). These prophages replicate passively with the host cell until specific stimuli induce the switch to the lytic replication cycle, resulting in bacterial lysis and the release of new virions (14). This induction is typically associated with the bacterial SOS response—an inducible DNA repair system regulated by the LexA repressor and the RecA inducer (15-17). While antibiotic exposure is a well-characterized trigger of the SOS pathway and subsequent prophage induction, host bacteria are often exposed to an extraordinary range of stressors and compounds that could be driving the microbial dynamics through the induction of prophages.

Previous studies have identified various foods and compounds capable of inducing prophages. For example, Oh *et al*. reported that *Lactobacillus reuteri* prophages can be induced with dietary fructose and short-chain fatty acids via RecA-dependent activation of the Ack pathway (18). Similarly, Silpe *et al*. demonstrated that colibactin, a genotoxin produced by certain gut microbiome members, triggers prophage induction in *Salmonella* via the SOS response (19). While these studies explored mechanistic links between specific compounds and phage activation, they focused on a narrow set of inducers.

However, broader screens have begun to emerge. For example, Boling *et al*. assessed 117 foods and found that compounds such as artificial sweeteners and bee propolis increased virus-like particle (VLP) counts and inhibited bacterial growth in a species-specific manner (20). Validation of selected hits was limited to VLP quantification by flow cytometry—a method that cannot distinguish between infectious and non-infectious particles. Similarly, Tompkins *et al*. screened 3,747 compounds using a high-throughput luminescent β-galactosidase assay which led to the identification of four-naturally-derived compounds as potential prophage inducers (21). However, the method used measures phage gene expression and not actual phage production or infectivity. Other studies have more broadly examined the effects of commonly consumed compounds on the gut microbiome. Sutcliffe *et. al*., tested 12 medications at five different concentrations against eight bacterial isolates; yet the methods of validation faced similar limitations to those described above (22).

These studies highlight that diverse environmental and dietary agents can modulate bacterial and phage dynamics. However, the extent to which frequently encountered compounds broadly influence prophage induction remains largely unexplored. Systematic, functionally validated approaches—particularly those confirming infectious phage production—are essential for uncovering overlooked interactions and expanding our current understanding of phage–host dynamics. Addressing this gap will provide researchers with new tools to broaden the study of prophage biology, enabling investigation into alternative induction pathways, particularly in cases where compounds are not traditionally associated with DNA damage. This is especially important given the field’s historical reliance on canonical inducers such as Mitomycin C and Ciprofloxacin, which may obscure other ecologically or clinically relevant triggers.

Here we employ High-Throughput Screening (HTS) using a large compound library to identify compounds capable of triggering prophage induction in model lambdoid prophage HK97 (23). When lysogenic bacteria are exposed to compounds capable of inducing the lytic cycle, phage-mediated lysis may occur, resulting in reduced endpoint optical density (OD) at concentrations that would not necessarily affect non-lysogenic counterparts (24). This differential growth phenotype provides a scalable and accessible readout for HTS. By comparing the growth dynamics of lysogenic and non-lysogenic strains, candidate inducers can be identified and subsequently validated through direct phage quantification. This approach enables systematic discovery of previously unrecognized prophage inducers and lays the groundwork for exploring alternative induction pathways beyond canonical DNA damage responses.

## Results and Discussion

### HTS identified several novel prophage inducers

We conducted a primary screen using *E. coli* K12 and its HK97 lysogen exposed to 3,921 bioactive compounds to identify compounds capable of inducing prophage-mediated bacterial killing. Ciprofloxacin at MIC (minimum inhibitory concentration) served as a positive control for growth inhibition and z’ factors were calculated to ensure assay quality (Table S1). : While media-only negative controls showed no growth and cell-only conditions exhibited no inhibition, the Ciprofloxacin MIC control resulted in growth inhibition of both strains, reducing growth to 18% (HK97 lysogen) and 20% (*E. coli* K12) relative to untreated controls, as expected. Because the ciprofloxacin control was added at MIC of the more resistant (non-lysogen) strain, it should not differentially affect the two strains despite the known inducing properties of the antibiotic.

Compounds were designated as primary hits if exposure of HK97 lysogen to the compound resulted in less than 50% bacterial growth relative to the untreated control on the same plate, and if growth of the HK97 lysogen was 70% or less compared to *E. coli* K12 treated with the same compound. These criteria were selected to enrich for compounds that cause increased bacterial killing in the presence of the prophage. Application of these thresholds yielded 37 primary hits (Figure 1a, Table S2). These compounds produced a greater killing effect in the presence of both the compound and the prophage, consistent with the sensitization we expect from prophage induction. A single compound, Benzoxyquine, resulted in a massive benefit to the lysogen – sitting at coordinates 97, 291, and was omitted from Fig 1a to increase legibility.

**Figure 1:**
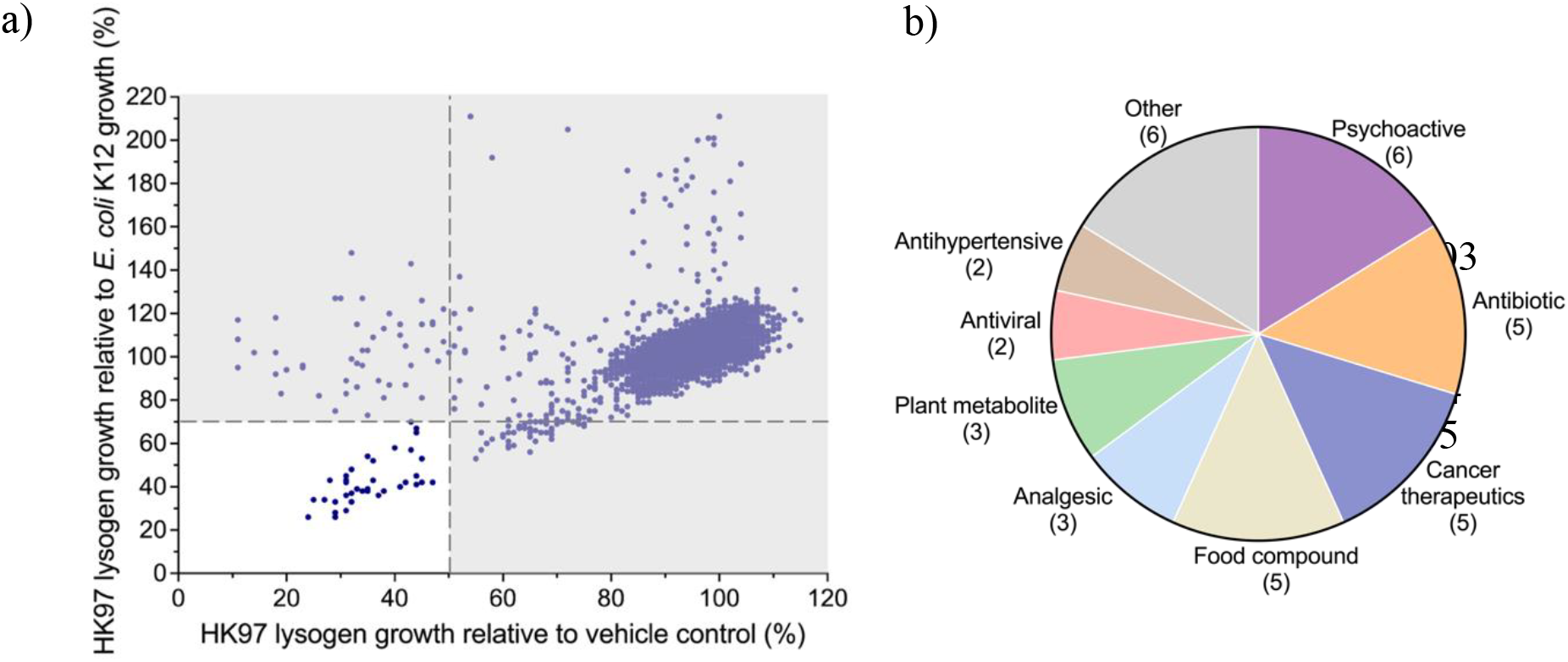
High-throughput screening identified 37 primary hits identified as potential inducers. (a) Primary hits identified in the HK97 lysogen model. Each dot represents a compound, and hits are defined as compounds that, averaged across both replicates, cause 50% or less growth relative to a no compound control and 70% or less growth relative to *E. coli* K12 + compound. These are represented in the white box. (b) Categorization of primary hits.

Notably, several known SOS pathway inducers such as fluoroquinolones, Lomefloxacin hydrochloride and Ofloxacin, were identified among the hits, lending support to the validity of the screen (25, 26). However, the identified compounds spanned a wide variety of pharmacological classes, including antivirals, antineoplastics, analgesics, and dietary supplements (Figure 1b), suggesting that selection was not restricted to traditional antibiotics. This diversity highlights the potential to identify novel prophage inducers. However, because hits were identified based on growth inhibition, which was used as a proxy for prophage induction, we carried out further validation to confirm whether these compounds directly trigger prophage induction.

To validate selected primary ‘hits’, 13 compounds were ordered for further analysis, based on a combination of cost and functional diversity: Valporic Acid, Thiothixene HCl, Chlorambucil, Chlorophyllide Cu complex Na salt, Astaxanthin, Gibberellic acid, Docosanol, Oxoantel Pamoate, Imiquimoid, Berberine, Fluoxetine HCl, Harmane, Bleomycin (Figure 2a). Of these 13, eight failed to yield any MIC in our larger volume assays. For two more, the MICs could not be determined due to compound precipitation and pigmentation. However, Berberine, Harmane, Fluoxetine HCl, and Bleomycin were verified as prophage inducers (Figure 2b).

**Figure 2:**
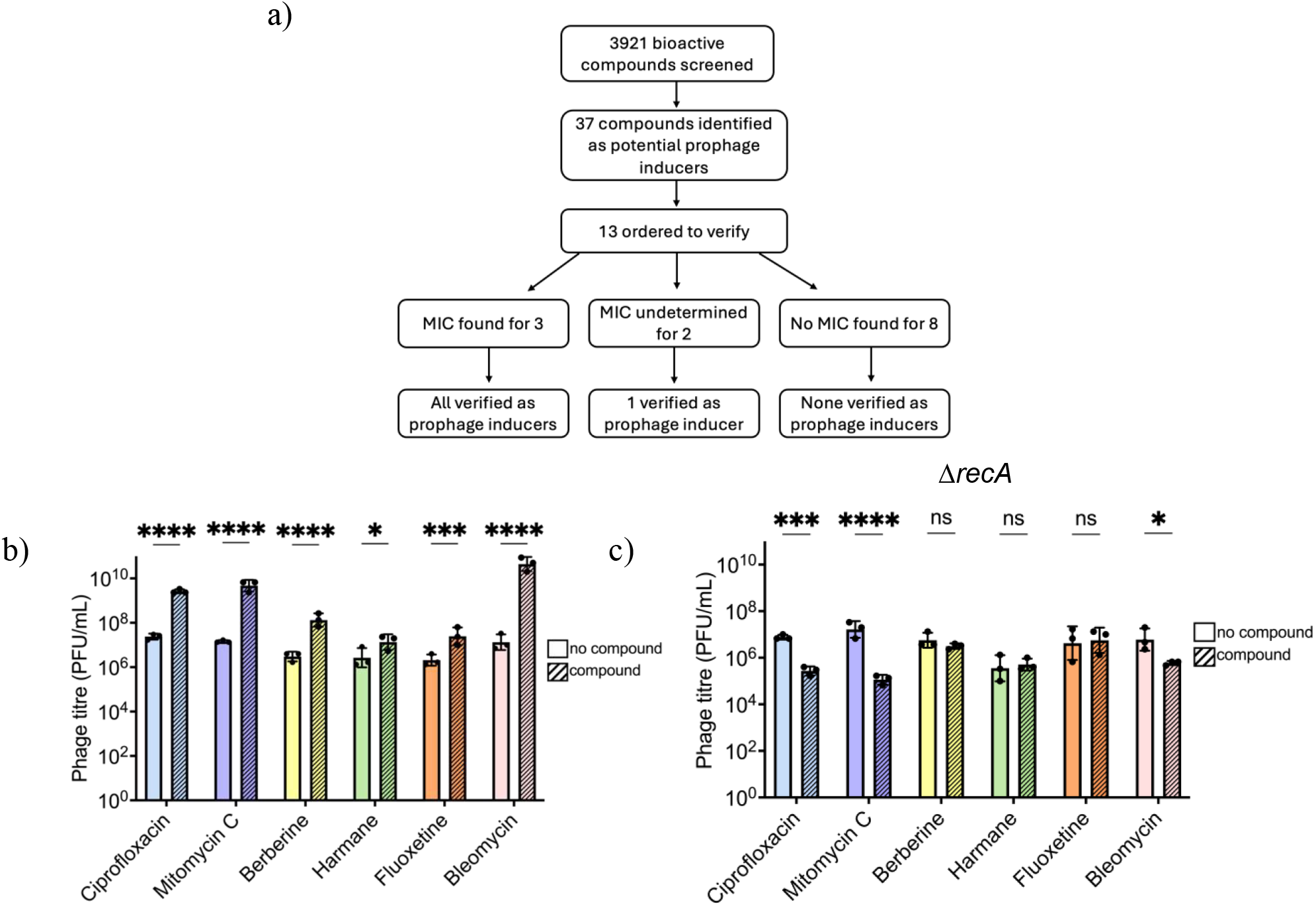
Characterized primary hits are SOS-dependent prophage inducers. (a) Summary of the validation process of ordered hits. MIC is defined as the minimal inhibitory concentration. (b) Phage quantification of HK97 lysogen filtrates alone and challenged with different compounds across three biological replicates. (c) Phage quantification of *ΔrecA* lysogen lysates alone and challenged with different compounds across three biological replicates. Error bars represent SD. Titres compared using two-way ANOVA and Šidák’s multiple comparison tests. * p ≤ 0. 05, ** p ≤ 0. 01, *** p ≤ 0. 001 and **** p ≤ 0.0001. ns = non significant.

Berberine, a plant alkaloid with weak antibiotic activity, is the primary bioactive component of tree turmeric and has been traditionally used in Ayurvedic and Chinese medicine (27). It has also been shown to modulate metabolic disorders, including type 2 diabetes and dyslipidemia, through AMPK activation and enhancement of glycolysis, leading to reduced insulin resistance (28, 29). MIC could not be determined, as compound pigmentation and precipitation interfered with spectrophotometric assays. Regardless, lysogen was subjected to a range of Berberine concentrations and the greatest magnitude of induction (∼2-log increase) relative to baseline spontaneous induction levels was observed at 350 μg/mL (Figure 2b).

Harmane, a heterocyclic β-carboline, is found in commonly consumed products such as cooked meats, coffee, soy sauce, and tobacco products (30). It functions as a monoamine oxidase inhibitor and has been linked to altered dopamine metabolism (31, 32). Harmane MIC for *E. coli* K12 was found to be 100 μg/mL and exposure of HK97 lysogen to ½ MIC resulted in approximately 1-log increase in phage titer relative to baseline spontaneous induction levels (Figure 2b).

Fluoxetine HCl (Prozac), a selective serotonin reuptake inhibitor (SSRI), is widely prescribed for the treatment of mood disorders and is among the most frequently used antidepressants in Canada, where approximately 17% of the population is prescribed antidepressants (33, 34). Fluoxetine MIC for *E. coli* K12 was found to be 100 μg/mL and exposure of HK97 lysogen to ½ MIC resulted in approximately 1-log increase in phage titer relative to baseline spontaneous induction levels (Figure 2b). Notably, Berberine, Fluoxetine, and Harmane have all been reported to impact the gut microbiome and/or exhibit antibacterial properties, suggesting a potential role for prophages in mediating their effects (35-37). Although none of these compounds are currently recognized as prophage inducers, recent evidence indicates that Fluoxetine can activate the bacterial SOS response in *E. coli*, resulting in Shiga Toxin production which stx phages are known to mediate (38, 39). These findings further emphasize the likelihood that induced phages may play an existing role in altering microbiome composition, influencing the therapeutic effects of these compounds.

Lastly, Bleomycin is a glycopeptide antibiotic primarily used as an antineoplastic agent to treat squamous cell cancer, Hodgkin’s lymphoma and testicular carcinoma (40). Bleomycin MIC for *E. coli* K12 was found to be 3.125 μg/mL and exposure of HK97 lysogen to ½ MIC resulted in a greater than 3-log increase in phage titer relative to baseline spontaneous induction levels (Figure 2b) exceeding induction levels observed with Ciprofloxacin and Mitomycin C. It had once previously been identified as a phage inducer, but this exceptionally high induction had not been remarked upon or delved into (41).

Induction by all four compounds was dependent on the SOS response, as no induction was observed in an HK97 *ΔrecA l*ysogen where the SOS pathway is disabled (Figure 2c). Together, these findings validate that the compounds for which MICs could be obtained are prophage inducers, thereby supporting the overall robustness of the screening strategy.

Physiological concentrations of these compounds vary depending on dosage, metabolic rate, time since administration, and route of delivery. For Berberine, a widely used dietary supplement, the recommended daily dose is approximately 1500 mg throughout the day, where peak plasma levels reach 0.4 ng/ml after a single dose of 400 mg, substantially lower than the concentrations used in our assays (42). However, due to its common use and lack of dosing standardization, local concentrations in the gastrointestinal tract may exceed recorded systemic levels. Harmane levels are difficult to estimate accurately, given its presence in a range of commonly consumed foods such as meats, coffee, and tobacco products. Fluoxetine, at a maximum recommended daily slow-release dose of 90 mg, reaches plasma concentrations around 91 - 302 ng/mL (0.091 – 0.302 μg/mL), also lower than our assay concentrations (43). In contrast, Bleomycin achieves plasma levels that can overlap with the concentrations used in our screen, depending on administration route, cancer type, and patient-specific factors (44).

Unlike the other compounds identified, Bleomycin is a cytotoxic antibiotic and is unlikely to be used intentionally to modulate the microbiome. However, the magnitude of prophage induction observed with Bleomycin exceeds that of Ciprofloxacin and Mitomycin C— both well-established, SOS-dependent inducers—highlighting its potential as a powerful tool for studying prophage induction.

### Characterization of Bleomycin as an alternative to standard prophage inducers

To further characterize Bleomycin activity, HK97 lysogen was exposed to a range of Bleomycin concentrations (400 μg/mL – 0.0003 μg/mL). For comparison, the same was done with Ciprofloxacin and Mitomycin C. Ciprofloxacin induced at least a 1-log increase in phage titres relative to baseline spontaneous induction at concentrations between 0.25 μg/mL – 0.004 μg/mL (Figure 3). Mitomycin C displayed broader and more potent activity with at least 1-log induction across concentrations 25 μg/mL – 0.002 μg/mL (Figure 3). Compared to Ciprofloxacin and Mitomycin C, Bleomycin induces at an overall higher magnitude across a broader concentration range with at least a 1-log induction at concentrations 200 μg/mL – 0.002 μg/mL (Figure 3). Despite the strong prophage induction observed with Bleomycin, this effect does not correlate with magnitude of SOS activation, measured by *recA* expression, which was higher in Ciprofloxacin (Figure S1). Interestingly, the broad range of Bleomycin concentrations that induced phage production corresponded with a relatively stable level of *recA* expression across the same tested concentrations.

**Figure 3:**
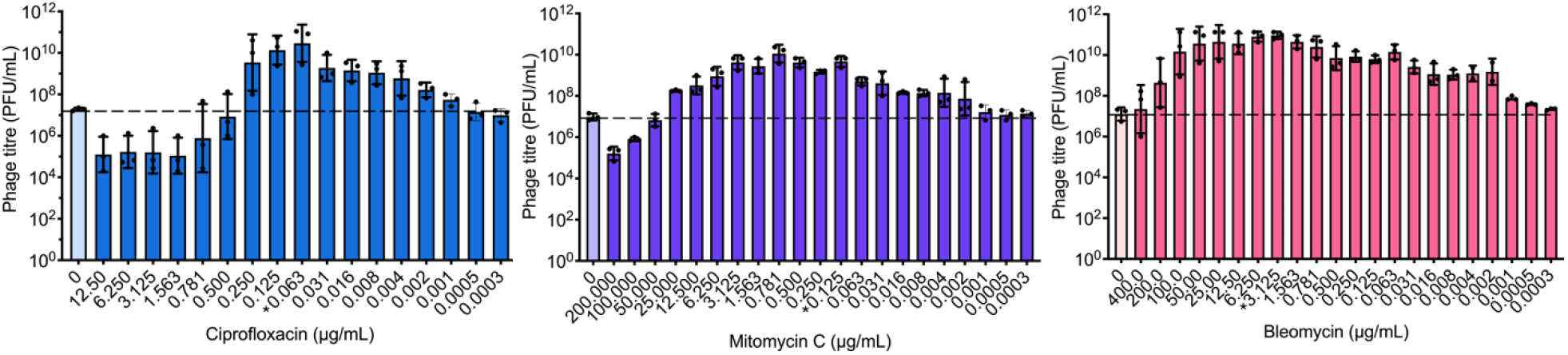
Bleomycin demonstrates greater activity compared to Ciprofloxacin and Mitomycin C. Phage quantification of HK97 lysogen either alone, or challenged with Ciprofloxacin, Mitomycin C or Bleomycin across a range of concentrations, averaged across 3 biological replicates. * Denotes MIC. Dotted line represents baseline induction levels. The bars represent the means, and the error bars the standard deviations.

Traditional inducers like Ciprofloxacin and Mitomycin C often require dose optimization and laborious troubleshooting due to their limited range and magnitude of activity. In contrast, Bleomycin’s broader effective concentration range and higher magnitude of induction reduces the need for optimization, serving as a more flexible alternative. To determine whether this represents a generalizable phenomenon, we tested Bleomycin’s ability to induce other phages, in other bacteria.

To investigate this, we exposed bacteriophage Mu lysogens to a range of Ciprofloxacin, Mitomycin C and Bleomycin concentrations. Mu is a temperate transposable phage that infects *E. coli* 40 by transposing its DNA into the host chromosome, almost at random (45). While no chemical treatments have previously been reported to induce Mu, induction has been achieved through specific viral mutants (46). Here, three Mu lysogens were used, each characterized by distinct integration sites confirmed via arbitrary PCR. Unexpectedly, exposure of Mu lysogens to Ciprofloxacin, Mitomycin C and Bleomycin were able to induce Mu in select lysogens. All three compounds were able to induce lysogen 3, but the effect was lost completely in lysogen 2 and partially in lysogen 1 (Figure 4a). This suggests that Mu is inducible under certain genomic contexts with host integration potentially influencing sensitivity.

**Figure 4:**
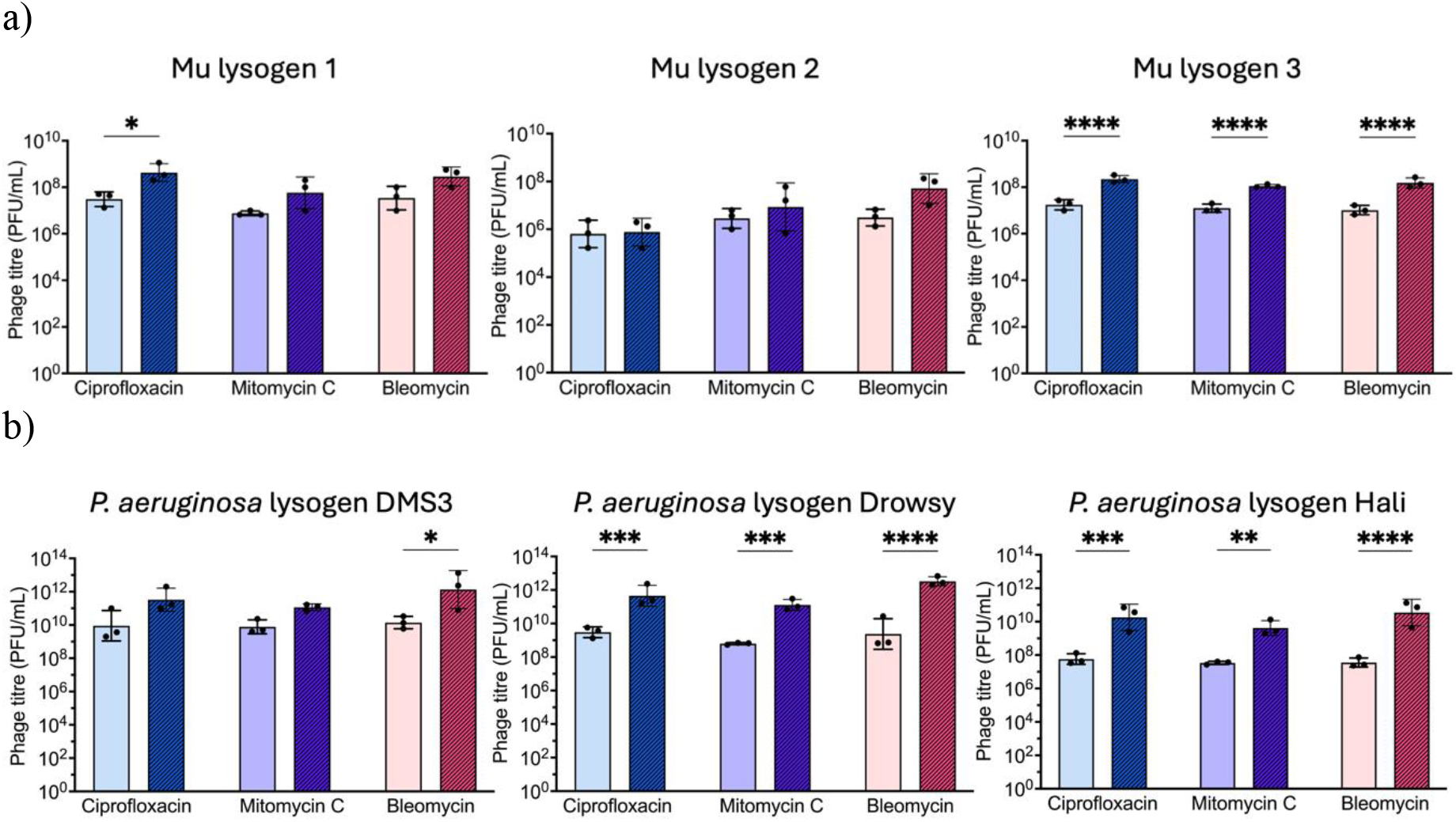
Effects of Bleomycin are generalizable across phages and bacterial hosts. (a) Phage quantification of *E. coli* phage Mu lysogen lysates alone and challenged with Ciprofloxacin, Mitomycin C or Bleomycin, averaged across three biological replicates. Lysogen number corresponds to the phage integrating at a different site within the host genome. (b) Phage quantification of different *P. aeruginosa* lysogen lysates alone and challenged with Ciprofloxacin, Mitomycin C or Bleomycin, averaged across three biological replicates. Error bars represent SD. Titres compared using two-way ANOVA and Šidák’s multiple comparison tests. * p ≤ 0. 05, ** p ≤ 0. 01, *** p ≤ 0. 001 and **** p ≤ 0.0001.

To determine if Bleomycin’s activity extended to other bacterial species, we tested its effects along with Ciprofloxacin and Mitomycin C on three *Pseudomonas aeruginosa* PA14 lysogens, each harboring Mu-like phages related to known *Pseudomonas* transposable phages (47). In contrast to the broader induction profiles seen with HK97, phage induction in *P. aeruginosa* was limited, with induction typically occurring at a single concentration for each compound. This occurred near the MIC for Ciprofloxacin of 0.063 μg/mL and near the MIC for Mitomycin C (0.125 μg/mL – 0.5 μg/mL). Bleomycin MIC was undeterminable with little reduction in O.D. at the highest concentration tested, yet induction was readily observed at 200 μg/mL. Ciprofloxacin and Mitomycin C induced only P8 and P13, while Bleomycin induced all three lysogens and resulted in higher magnitudes of induction, further emphasizing the compound’s potency and generalizability across different bacterial host and their prophages.

## Conclusions

This study employed HTS to identify several novel, commonly consumed compounds as prophage inducers, reinforcing the concept that phage induction is a key mechanism by which external agents can shape microbial community dynamics. Although most compounds were tested at supra-physiological concentrations, this does not preclude their capacity to act as prophage inducers, especially in the gut where localized exposure may be higher or more prolonged. The identification of widely used therapeutics and dietary supplements as inducers supports growing evidence that non-antibiotic agents can inadvertently modulate microbial ecosystems through phage-mediated effects (18-22). Notably, all validated inducers operated via the canonical SOS response, underscoring the conserved nature of this pathway.

Among these compounds, Bleomycin emerged as a potent and broadly effective prophage inducer, often outperforming current standard inducers Ciprofloxacin and Mitomycin C (6, 7). In addition to this performance, Bleomycin is also more cost effective and chemically stable compared to Mitomycin C, making it well-suited for routine laboratory use (48). Together these findings highlight the power of high-throughput approaches in identifying prophage inducers and establishes Bleomycin as a valuable tool for prophage research with the potential to replace currently used prophage inducers.

**Table 1.** Bacterial and Viral strains used in this study.

| Strain | Source |
| --- | --- |
| *E. coli* K12 HER1382 | Félix d’Hérelle Reference Center for Bacterial Viruses |
| *E. coli* K12 HK97 lysogen | Al-Anany *et. al.*, 2021<sup>49</sup> |
| *E. coli* BW25113 $\Delta recA$ | Dharmacon KEIO collection through Horizon Discovery (Cambridge, UK) |
| *E. coli* BW25113 $\Delta recA$ lysogen | This work |
| *E. coli* 40 | Félix d'Hérelle Reference Center for Bacterial Viruses |
| Mu lysogens | This work |
| *P. aeruginosa* PA14 | Burrows Lab, McMaster University |
| *P. aeruginosa* PA14 lysogen P8 | Fatima *et. al.*, 2025<sup>47</sup> |
| *P. aeruginosa* PA14 lysogen DMS3 | Fatima *et. al.*, 2025<sup>47</sup> |
| *P. aeruginosa* PA14 lysogen P13 | Fatima *et. al.*, 2025<sup>47</sup> |
| *E. coli* K12 MG1655 | Shao *et. al.*, 1997<sup>50</sup> |
| *E. coli* K12 MG1655 *recA* - GFP | Zaslaver *et. al.*, 2006<sup>51</sup> |
| *E. coli* K12 MG1655 *sulA* - GFP | Zaslaver *et. al.*, 2006<sup>51</sup> |
| *E. coli* K12 MG1655 *pitB* - GFP | Zaslaver *et. al.*, 2006<sup>51</sup> |

All bacterial strains are grown at 37°C shaking at 130 rpm or 250 rpm in LB broth.

## Methods

### Bacterial strains

### Generating HK97 Δ*recA* lysogen

To generate the HK97 Δ*recA* lysogen, serial dilutions of HK97 phage were plated on a soft agar overlay (10 mL 1% LB + 3 mL 0.75% LB with 300 μl of culture) of Δ*recA* and incubated overnight at 37°C. Regrowth was re-streaked, incubated overnight and 32-48 colonies were picked and purified. Each purified colony was then streaked on top of a soft overlay containing parental (phage sensitive) Δ*recA* strain, using a HK97 lysogen as a positive control. Plates were incubated overnight, and HK97 Δ*recA* lysogens were identified by the presence of zones of clearing surrounding the streak.

HK97 lysogen confirmation was done using PCR primers attBF and HK97lysR detecting the known phage-host junctions. The absence and presence of *recA* gene was verified by PCR using primers binding both within (RecA_Fwd and RecA_Rev) and outside (RecA_F and RecA_R) the gene, and further validated by whole-genome sequencing.

**Table 2.**
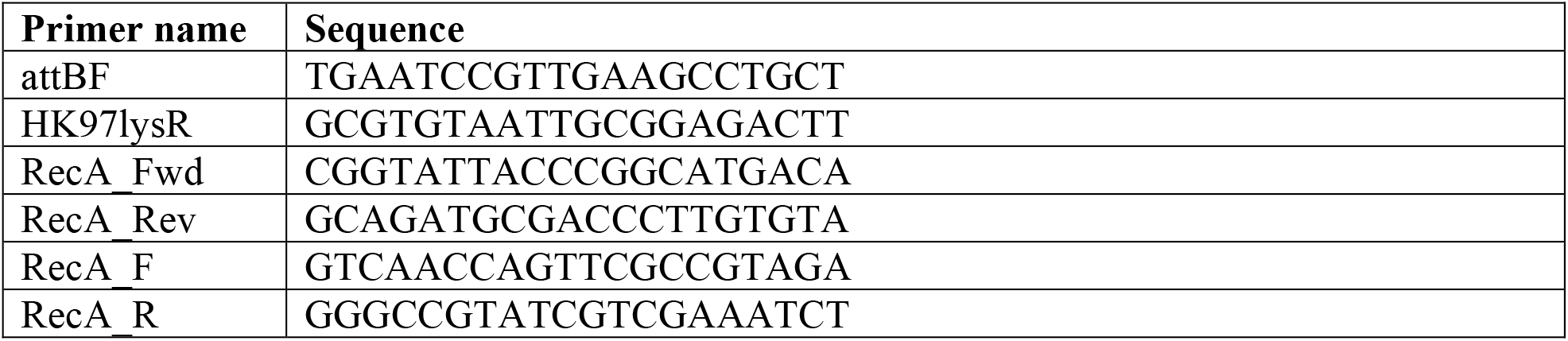
Primers used in this study.

### Prophage inductions assays

Induction assays were performed in 96-well plates using two-fold serial dilutions of the compound and incubated at 37 °C for 18 h. Optical density (O.D.) readings were taken either every 30 minutes or only at the endpoint (18 h). Cultures were added at an initial O.D. of 0.2, prepared by a 1:100 sub-inoculation from an overnight culture. Each well had a final volume of 250 μl. To confirm and test inducibility of lysogens, Ciprofloxacin and Mitomycin C were used as positive controls. MIC and ½ MIC of compounds were determined using the parent bacterial host and applied to corresponding lysogens. When the MIC could not be determined, induction assays were done using the highest compound concentration permissible, considering solubility and the percentage of organic solvent used. Compounds were either dissolved in either PCR H_2_O or dimethyl sulfoxide (DMSO). In instances where DMSO was used, vehicle controls confirmed the solvent did not have any effect on phage titre (Figure S2).

**Table 3.**
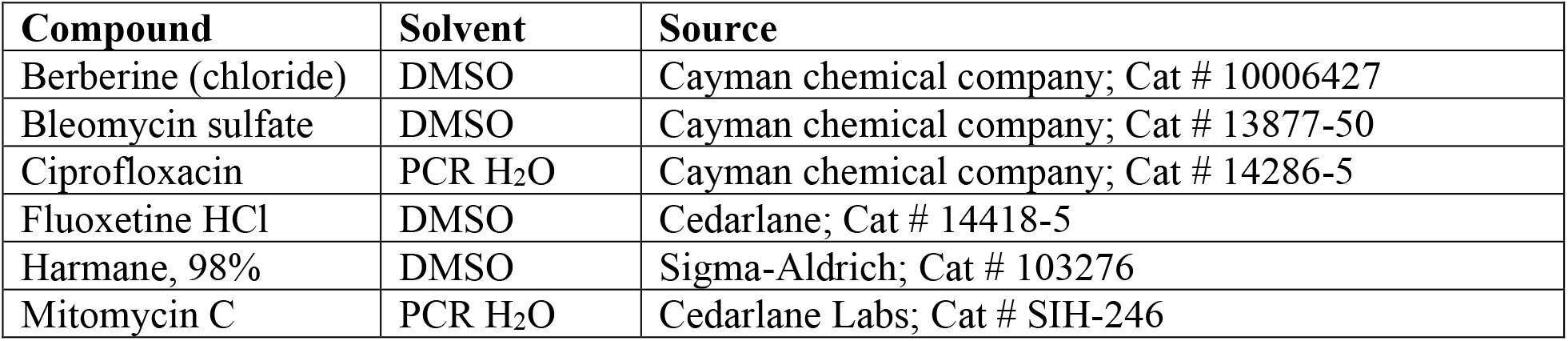
Compounds used in this study.

| Compound | Solvent | Source |
| --- | --- | --- |
| Berberine (chloride) | DMSO | Cayman chemical company; Cat # 10006427 |
| Bleomycin sulfate | DMSO | Cayman chemical company; Cat # 13877-50 |
| Ciprofloxacin | PCR H<sub>2</sub>O | Cayman chemical company; Cat # 14286-5 |
| Fluoxetine HCl | DMSO | Cedarlane; Cat # 14418-5 |
| Harmane, 98% | DMSO | Sigma-Aldrich; Cat # 103276 |
| Mitomycin C | PCR H<sub>2</sub>O | Cedarlane Labs; Cat # SIH-246 |

After incubation, plates were either filtered (MilliporeSigma Multiscreen High Volume Filter Plates - 0.45 μM) or centrifuged (2103 RCF, 25 min). Approximately 100 μl of the resulting supernatant was aspirated, serially diluted (10-fold) in LB broth, and plated (3 μl) on respective bacterial hosts (*E. coli* K12 Δ*recA, E. coli* K12, *E. coli* 40, PA14). When working with PA14 bacterial strains, biofilms were removed prior to centrifuging by inserting filter tips into the wells and allowing the biofilm to adhere. Phage concentration is calculated via the following formula:

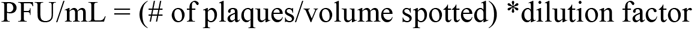

### High throughput screening

The HTS was an integrated system with most operations occurring offline, requiring little manual assistance. Each assay plate was barcoded prior to the experiment.

A total of 3921 of bioactive compounds from the Centre for Microbial Chemical Biology (CMCB, McMaster University, Hamilton, Ontario, Canada) collection were tested against two bacterial strains (*E. coli* K12 and *E. coli* HK97 lysogen) using 384-well plates. Each assay plate contained one bacterial strain grown to an O.D. of 0.4. The HTS pipeline involved dispensing the bioactive compound at a final concentration of 10 μM using the Echo Acoustic Liquid Handler (Beckman Coulter, California, USA), followed by the addition of controls—DMSO (negative control) and ciprofloxacin (positive control)—using the Multidrop Combi Reagent Dispenser (Thermo Fisher Scientific, Massachusetts, USA). Subsequently, 30 μl of culture was added using the Tempest Liquid Dispenser (Formulatrix, Dubai, UAE). Plates were incubated at 37 °C for 6 hours, with O.D. readings taken every 30 minutes using the Biotek Epoch Plate Reader (Agilent, California, USA). Bioactive compounds were added in duplicate at a final concentration of 10 μM with 30 μl of culture and 0.3 μl of compound added. Each assay plate included a column of DMSO (1%) (compounds are diluted with DMSO) + LB media (cells are grown in LB), a column of DMSO (1%) + MIC Ciprofloxacin + LB media, a column of DMSO (1%) + cells and a column of DMSO (1%) + MIC Ciprofloxacin + cells. These conditions served as negative and positive controls. Screening was performed for a total of 6 h, with reads taken every 30 min.

#### Data analysis

Percent growth relative to the average of each plate’s untreated controls was calculated individually for each plate and then averaged across the two technical replicates. O.D_600_ readings obtained at 6 h were used for analysis.

### SOS gene expression assay

To quantify the magnitude of SOS pathway activation, three *E. coli* strains containing engineered promoter-reporter gene constructs were used, each expressing GFP in response to activation of *recA, sulA*, or *pitB* expression. Assays were done in 96-well plates with two-fold dilutions of compound added to 100 μL of culture grown to O.D. 0.2. Plates were incubated at 37°C for 18 h, after which absorbance and fluorescence reads (479, 520) were obtained. Strains were maintained in the presence of Kanamycin (50 μg/mL) in overnight cultures to ensure plasmid selection. Resulting fluorescence values are normalized to O.D., then further normalized to the fold change observed in *pitB* expression. *E. coli* MG1655 (WT) was used as the negative untreated control. Assays were done in biological duplicates, each with technical duplicates.

### Statistical Analysis

All figures and statistical analyses were generated using GraphPad Prism version 10.3.1 (GraphPad Software, Inc., CA, USA). Flowcharts were created using Microsoft PowerPoint.

## Supplementary

**Figure S1:**
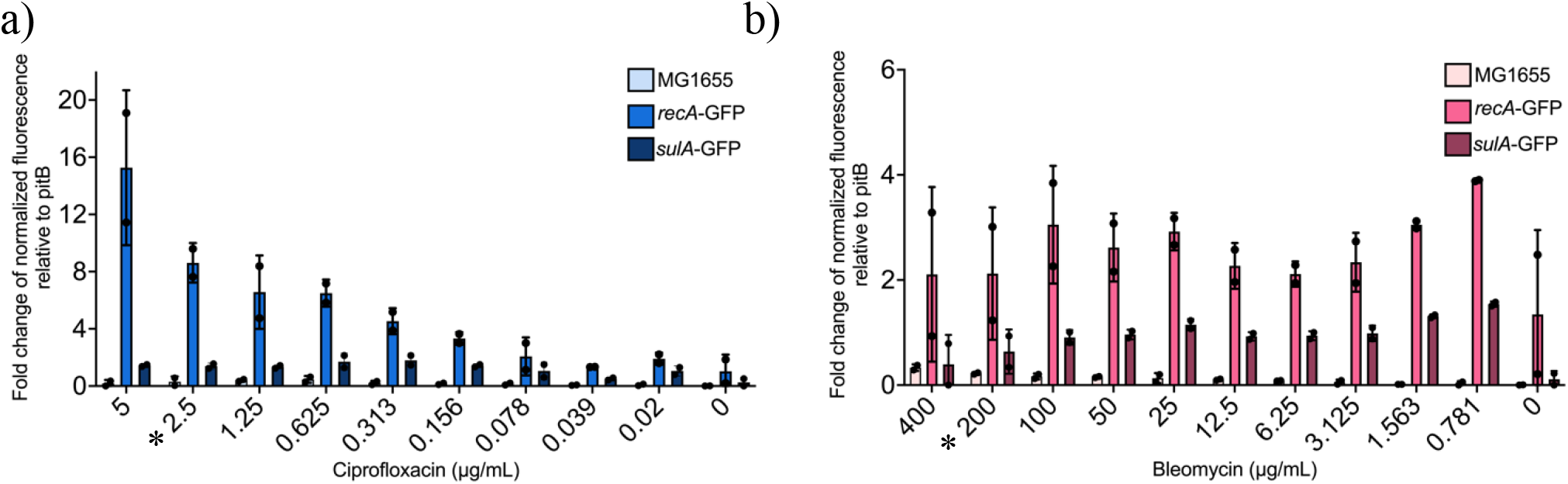
Exposure to Bleomycin results in consistent levels of SOS induction across a concentration range. (a) Magnitude of SOS induction when strains are exposed to a range of Ciprofloxacin concentrations averaged across two biological replicates. (b) Magnitude of SOS induction when strains are exposed to a range of Bleomycin concentrations averaged across two biological replicates. * denotes MIC. Error bars represent SD.

**Figure S2:**
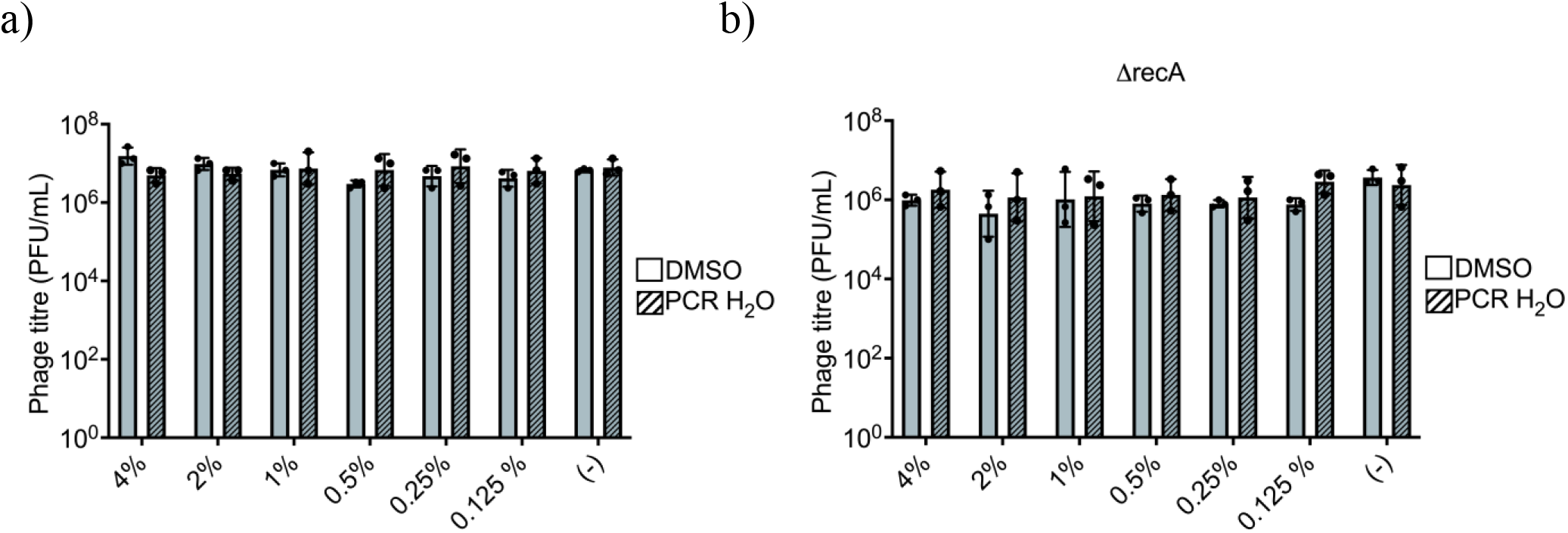
Solvent used to dissolve bioactive compounds have no effect on induction. (a) Phage quantification of HK97 lysogen alone and challenged with DMSO or PCR H_2_O averaged across three biological replicates. (b) Phage quantification of HK97 Δ*recA* lysogen alone and challenged with DMSO or PCR H_2_O averaged across three biological replicates. Error bars represent SD.

**Table S1:** Z’ Factor Determination for HTS Assay Validation.

| Time point | *E. coli* K12 + 1% DMSO | *E. coli* HK97 lysogen + 1% DMSO |
| --- | --- | --- |
| 5 hrs | 0.7868 | 0.6585 |
| 5.5 hrs | 0.8130 | 0.6681 |
| 6 hrs | 0.8085 | 0.6823 |

**Table S2:** List of primary bioactive compounds hits. Organized from greatest effect on OD to least effect. Highlighted cells indicate validated prophage inducers.

|  | Bioactive Compound |
| --- | --- |
| 1 | Meptazinol hydrochloride |
| 2 | Desoxymetasone |
| 3 | Alanyl-dl-leucine |
| 4 | Valproic acid sodium |
| 5 | Valacyclovir Hydrochloride |
| 6 | Harmane |
| 7 | Epirubicin Hydrochloride |
| 8 | Tolazoline Hydrochloride |
| 9 | Chlorophyllide Cu Complex Na salt |
| 10 | 1,3-Dipropyl-8-p-sulfophenylxanthine |
| 11 | Thiothixene hydrochloride |
| 12 | Lercanidipine hydrochloride hemihydrate |
| 13 | Olomoucine |
| 14 | Chlorambucil |
| 15 | Tramadol Hydrochloride |
| 16 | Oxantel pamoate |
| 17 | Lomefloxacin hydrochloride |
| 18 | AL-8810 |
| 19 | Docosanol |
| 20 | Imiquimod |
| 21 | Betulinic acid |
| 22 | Lobaric acid |
| 23 | Ethyl Everninate |
| 24 | Gibberellic acid |
| 25 | Willardiine |
| 26 | Salidroside |
| 27 | Liquiritigenin Dimethyl Ether |
| 28 | Lycorine |
| 29 | Ofloxacin |
| 30 | Astaxanthin |
| 31 | Isoquinoline hydrochloride |
| 32 | Berberine |
| 33 | Nystatine |
| 34 | Fluoxetine hydrochloride |
| 35 | Bleomycin B2 |
| 36 | Rolitetracycline |
| 37 | Cefepime hydrochloride |

